# Natural DEET substitutes that are strong olfactory repellents of mosquitoes and flies

**DOI:** 10.1101/060178

**Authors:** Sean Michael Boyle, Tom Guda, Christine Krause Pham, Sana Khalid Tharadra, Anupama Dahanukar, Anandasankar Ray

**Author notes:** **CORRESPONDING AUTHOR**: Anandasankar Ray Department of Entomology, University of California Riverside, 3401 Watkins Drive, Riverside, CA 92521, USA.

## Abstract

Despite shortcomings, N,N-Diethyl-3-methylbenzamide (DEET) has remained the gold-standard of insect repellents for >60 years. There are significant impediments to finding improved substitutes because the molecular targets causing repellency are unclear, new chemistries will require significant human-safety testing, and predicted costs for development are exorbitant. Here we identify shared structural features important for repellency and using a supervised chemical-informatics method screen *insilico* >400,000 compounds to identify >100 natural compounds as candidate repellents. We select 4 candidates that are affordable, 3 approved as safe for human food use, and demonstrate that they are strong olfactory and gustatory repellents to both mosquitoes and *Drosophila*. The chemicals do not dissolve plastic and have a mild and pleasant odor. These repellents are representative of a new generation of affordable substitutes for DEET that can be rapidly deployed globally because of excellent human-safety profiles, and have great potential in reducing deadly diseases by reducing mosquito-human contact.

## INTRODUCTION

Blood-feeding insects transmit deadly diseases such as malaria, Dengue, Filariasis, West Nile Fever, Yellow fever, Sleeping sickness and Leishmaniasis to hundreds of millions of people, causing untold suffering and more than a million deaths every year. Insect repellents can be very effective in reducing vectorial capacity by blocking the contact between blood-seeking insects and humans, however they are seldom used in disease-prone areas of Africa and Asia due to high costs and need for continuous application on skin.

*N*,*N*-Diethyl-m-toluamide (DEET) has remained the primary insect repellent used for more than 60 years in the developed world. DEET is a solvent, melting several forms of plastics, synthetic fabrics, painted and varnished surfaces(Krajick, 2006). Additionally, DEET has been shown to inhibit mammalian cation channels and human acetylcholinesterase, which is also inhibited by carbamate insecticides commonly used in disease endemic areas (Corbel et al.,2009). These concerns are enhanced by the requirement of direct and continuous application of DEET to every part of exposed skin in concentrations that can be as high as 30-100 percent. Several instances of increased resistance to DEET have also been reported in flies (Reeder et al., 2001), *Anopheles albimanus*(Klun et al., 2004), and *Aedes aegypti* (Stanczyk et al., 2010). Moreover, mosquito strains with resistance to pyrethroid insecticides, the main line of defense against mosquitoes in developing countries, are spreading (Butler, 2011). There is therefore an urgent need to develop safer and improved alternatives to DEET. The other major barrier in developing new repellents is the time and cost of development. It has been suggested that >$30M and several years may be required for identification (Gupta, 2007) and adequate human-safety analysis of new repellent chemistries.

A major limitation to finding effective substitutes is that the molecular targets in adult mosquitoes through which DEET causes repellence are unknown. Here, we describe identification of novel repellent odors that are safe, inexpensive, and effective at repelling *D. melanogaster* and *A. aegypti*. We first demonstrate that *D. melanogaster* and *A. aegypti* may utilize their gustatory system and olfactory system to different extents to avoid DEET, with mosquitoes relying more heavily on olfactory pathways. We then use a novel *chemical informatics* approach that uses supervised training from known repellents to identify important structural features that are responsible for avoidance behavior. Using these features we predict novel repellents from a very large untested odor space comprising a large purchasable odor set and a natural odor library, the latter providing many chemicals that are safe and already approved for human use by both United States and European governmental safety organizations. We select four odors from the 150 natural odor library predictions and demonstrate that all four are able to cause strong avoidance in both *D. melanogaster* and *A. aegypti*. Due to the large number of candidates we are able to select new repellents with ideal properties; safe for human consumption, do not dissolve plastics and mild and pleasant aroma. These results suggest that our integrative approach of computational predictions and behavioral analysis can revolutionize the discovery of safe and effective repellent odors that could be very useful in our struggle against the increasing spread of insect disease vector species. Although it has been several years since odor receptors were identified in vertebrates and invertebrates, very few odor receptor targets have been identified for known behavior modifying odorants. This study therefore has broader implications since the approach presented can be used for identification of improved behavior modifying odorants for any organism, even if the target odor receptor is unknown. Moreover upon identification of target receptors, the same methodology can be easily adapted for a receptor-activity based approach.

## RESULTS AND DISCUSSION

### *Aedes aegypti* Detect and Avoid DEET Primarily using Olfaction

To test whether the gustatory system of mosquitoes play a substantial role in DEET repellency as well, we developed a modified hand-in-glove assay (described in Supplementary Methods) (Figures 1A, 1B, and S1). The assay allows us to quantitatively analyze the repellent effects of DEET on mosquitoes attracted to a human arm, without being able to bite. Female *Aedes aegypti* mosquitoes show an equally strong avoidance behavior to DEET in both the contact and non-contact versions of the assay (Figure 1B). For rare landings, the time spent on the net before escape is lower, but not significantly different, when direct contact with DEET was permitted (Figure 1C). These results indicate that the female *Aedes* may use more of the olfactory system than gustatory to sense DEET directly and avoid it. This is consistent with previous observations that *Culex quinquefasciatus* (Syed and Leal, 2008) and *A. aegypti* (Turner et al., 2011) avoid DEET presented as a volatile stimulus directly. A DEET-sensitive neuron type has been identified in *C. quinquefasciatus* (Syed and Leal, 2008) and *A. aegypti* (Stanczyk et al., 2010), however it is not known whether these neurons are responsible for repellency, or which odor receptors they express. A broadly tuned larval odor receptor AgOr40 has been shown to respond to DEET (Liu et al., 2010; Xia et al., 2008), however its role in avoidance behavior in adults has not been determined.

**Figure 1.**
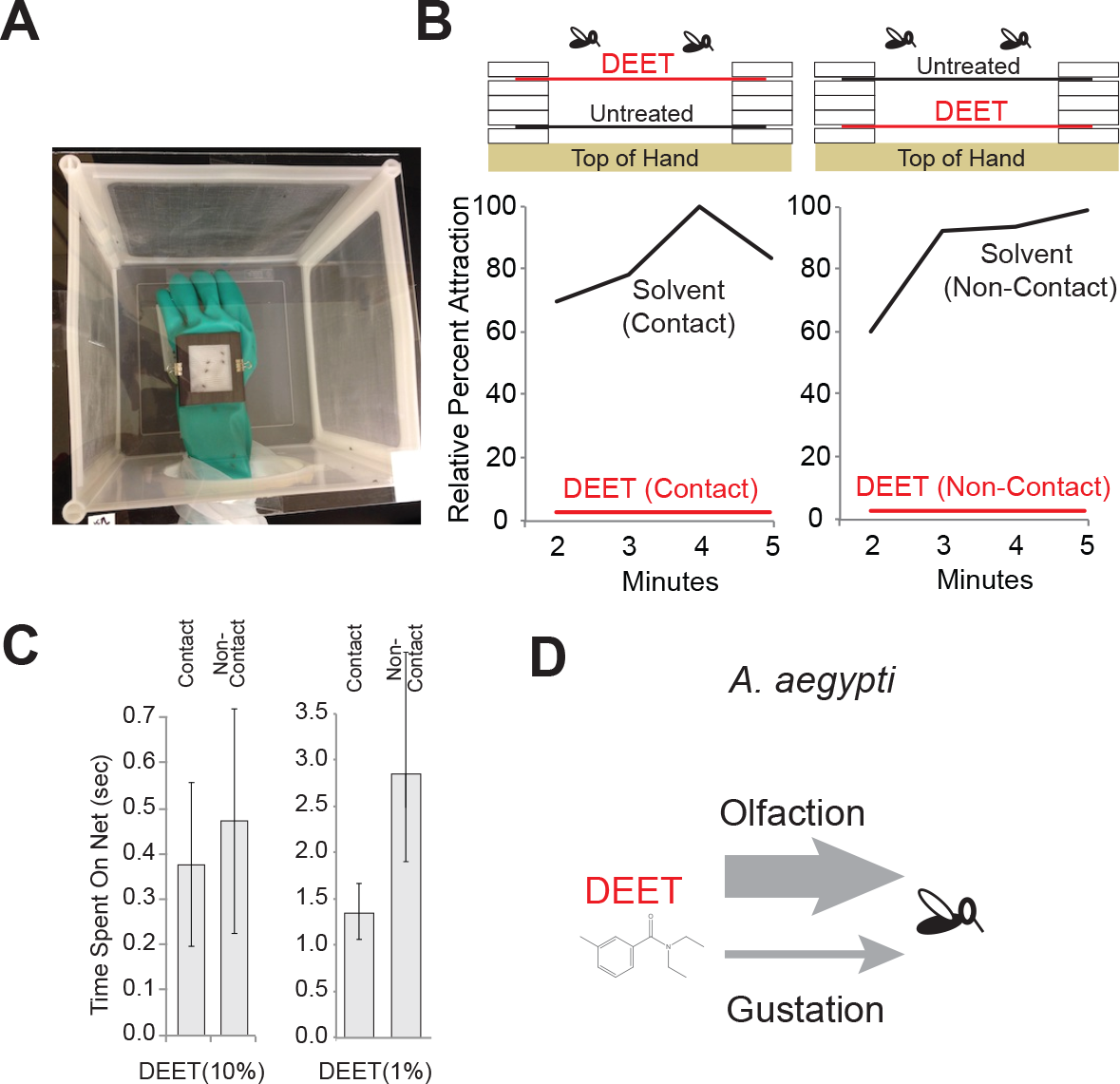
Contribution of olfaction and gustation in DEET avoidance. **A,** Photograph of the hand-in-glove assay to measure repellency in mosquitoes. **B,** Relative attraction of female *Aedes aegypti* in hand-in-glove assay measured as a percentage of mosquitoes spending >5-sec on the net covered window of glove, measured at 1 min intervals. (Left) DEET (10%) treated netting placed at the top level allowing contact, or (Right) at the lower level not allowing contact. N=5 trials for each, 10 mosquitoes/trial. **C,** Average time spent per mosquito on net for each landing event. **D,** Schematic of DEET mediated repellency in *Aedes aegypti*.

### A Computer can be trained to Predict Repellent Behavior from chemical structure

Volatile chemical space that can be exploited to find improved DEET substitutes is vast (potentially >400,000) thus using behavior assays alone is unfeasible from the perspective of time and cost of chemical purchase. Moreover, since a protein target for DEET action is unknown, high-throughput screening, or sophisticated computational protein-ligand docking based approaches to identify new ligands are also not possible.

To circumvent these problems so as to enable selection of compounds for behavior assays, we developed a novel chemical informatics approach. We hypothesized that the unknown target protein is recognizing specific structural features of DEET and other known structurally related repellents. By identifying structural features that are shared amongst DEET and the known repellent compounds, we can utilize them to rapidly screen an extremely large number of compounds *in-silico* to identify novel repellents, thus greatly reducing both the cost and time required. We assembled a training set of known repellents that included: the two approved ones DEET (a carboxamide) and picaridin; and also 34 N-acyl piperidiens(Katritzky et al., 2008) that are structurally related to picaridin; eucalyptol, linalool, alpha-thujone, beta-thujone(Kline et al., 2003; Klocke et al., 1987; Syed and Leal, 2008) and a structurally diverse panels of odors that are not expected to elicit repellency via similar target receptors(Carey et al., 2010; Hallem and Carlson, 2006). The study where the 34 n-acyl piperidiens were identified also showed that a chemical-structure-based approach could be successfully applied to predicting repellency(Katritzky et al., 2008). For our analysis, compounds from different sources were approximated into a single metric of “protection duration” as a rough indicator of repellency. The non-repellent diversifying training set of odors were assigned protection times of zero, while the approved repellents DEET and Picaridin were assigned the highest value since these would have the properties important for regulatory approval. Compounds were clustered using Euclidean distance and hierarchical clustering based on differences in repellency values, and a set of 5 compounds with the highest activity that clustered together was classified as “training repellents”.

We expect that only specific structural features of the repellent odors will interact with target proteins to elicit repellent responses, and not the entire molecule. We assumed that identification of structural features that are shared across repellent odors would enable a search for these features within a large chemical space, potentially identifying novel repellents (Maldonado et al., 2006). We decided to focus on a descriptor-based computational approach that is effective in structure analysis and is efficient in subsequently searching a large chemical space rapidly. We calculated mathematical values for 3,242 molecular descriptors, that describe the 3-D structure of a chemical, for our 201 compound training set and using a Sequential-Forward-Selection method(Haddad et al., 2008) we incrementally identified a unique subset of 18 descriptors that were highly correlated with repellency (correlation of 0.912)(Figures 2A, 2B, and 2C). As expected the repellent odors cluster together in the training set if the optimized descriptor subset is used to calculate Euclidean distance amongst them (Figures 2B). These 18 molecular descriptors represent a collection of predominately 2D and 3D descriptor types. Inspection suggests that 6 member rings, carbon-nitrogen distances, tertiary amides, and oxygen placement are prominent in the optimized subset. Interestingly, although repellents are topically applied chemicals, the Ghose-Viswanadhan-Wendoloski drug-like index is selected, which is an aggregate descriptor that usually suggests similarity of chemical features important for drugs (Ghose et al., 1999).

**Figure 2.**
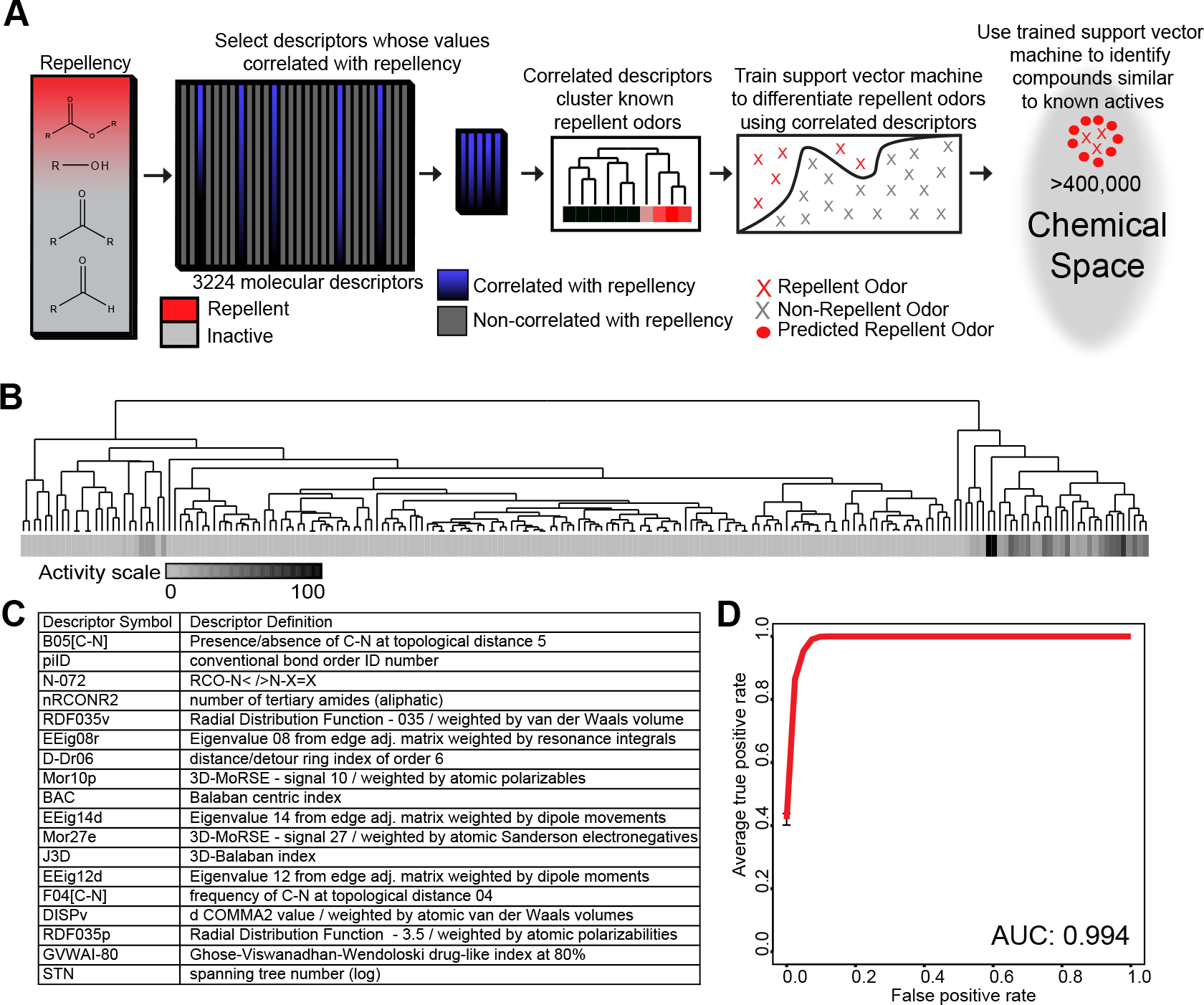
A chemical informatics method to predict repellency. **A,** Overview of the cheminformatics pipeline used to identify novel DEET-like ligands from a larger chemical space. **B,** Hierarchical cluster analysis of the 201 odorants of the training set using the optimized descriptor set to calculate distances in chemical space. **C,** Repellency-optimized descriptor symbols and brief descriptions arranged according to order in which they were selected for the optimized set. **D,** Receiver-operating-characteristic curve (ROC) representing computational validation of repellent predictive ability. The mean true-positive value from 20 independently run 5fold cross validations is plotted, where ~20% of the dataset was left out of training-set as a test-set for each run. The mean area under the curve (AUC) is provided.

In order to improve the predictive ability of the chemical informatics approach we used the optimized descriptor set to train a Support Vector Machine (SVM), which is a well-known supervised learning approach (Cortes and Vapnik, 1995)(Figures 2A). To validate the predictive ability of our approach, we performed 5-fold cross-validation using SVMs on the training set. Each cross-validation run excluded ~20% of the repellents as a test-set, while the remaining repellents were used to train a SVM. We predicted repellency for the withheld test-set odors using the trained SVM. This operation was repeated 5 times, each trial performed excluding a different subset of the training set, and the whole process was repeated 20 times for consistency. A mean Receiver-Operating-Characteristic (ROC) analysis curve representing the prediction accuracy was generated, and the Area-Under-Curve (AUC) value was determined to be 0.994, indicating that the *in-silico* approach was extremely effective in predicting repellents from compounds that are excluded from the training set (Figures 2D).

### High-Throughput *In Silico* Screen identifies Safe Natural Odors as repellents

We use the 18 optimized-descriptors and SVM method to screen *in-silico* a large chemical library consisting of >440,000 chemicals from a database called eMolecules of putative volatiles. Upon inspection, we find the top 1,000 predicted compounds (0.23% of hits) represent a structurally diverse group of chemicals that retain some structural features of the known repellents (Figures 3A and 3B). We computed logP values of the 1,000 compounds to identify ones that are predicted to be lipophilic (logP >4.5) thus allowing for selection of other compounds that are less likely to pass through the skin barrier in topical applications (Walker et al., 2003) (Figures 3B). In addition, we computed the predicted vapor pressures of these chemicals since volatility may predict ability of long-term protection vs. increased spatial domain of action (Figures 3B). Taken together the results of the screen present a very large collection of novel predicted repellents with desirable properties, identified via a computationally guided search of odor space.

**Figure 3.**
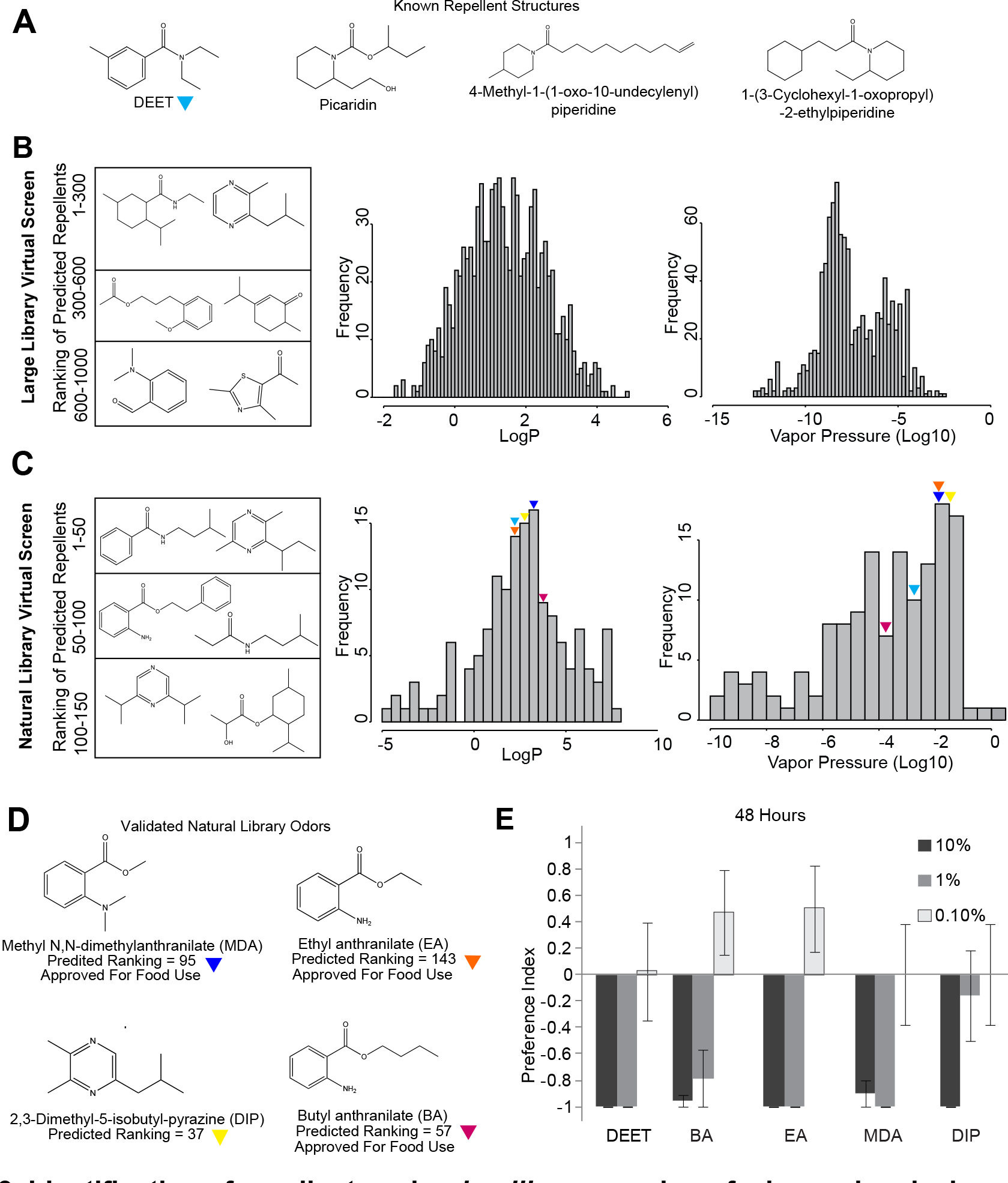
Identification of repellents using *in-silico* screening of a large chemical space. **A,** Examples of two approved repellents DEET and Picaridin, and two unapproved repellents (Katritzky et al., 2008). **B,** (Left) Representative predicted repellent odors from the odor library of >400,000. Computationally determined (Middle) LogP values and (Right) vapor pressure values for the top ranked 1000 predicted repellent compounds. **C,** (Left) Representative structures from the top 150 predicted repellent compounds from the natural odor library of >3,000. Computationally determined (Middle) LogP values and (Right) vapour pressure values for the top ranked 150 predicted repellent compounds. Color arrowheads indicate values for DEET and odors selected for behavior experiments from the natural library indicated in **D**. **E,** Preference index of Drosophila adults to predicted repellents at three different concentrations in a two choice trap assay measured after 48 hrs. N=7-10 trials each treatment (trials with <40% participation were excluded), 10 flies/trial, error bars=s.e.m

Although the *in-silico* screen bypasses the challenge of not knowing the protein target, the most significant challenge lies in identifying effective repellent substitutes for DEET that are affordable and safe and that can be rapidly approved for human use. In order to identify compounds that fit these criteria, we applied our *in-silico* screen to an assembled natural odor library consisting of >3,000 chemicals identified as either originating from plants, insects, or vertebrate species or compounds already approved for human use as fragrances, cosmetics or flavors (Supplemental Material). Using the trained SVM and optimized descriptor set on the natural library, we identified the top 150 ranked predicted repellent compounds. Predicted repellents share similarity in some parts of the structure while providing a diverse set of compounds (Figure 3C). For example, several anthranilates and pyrazines were identified that represent novel groups of safe and natural compounds, although such compounds were absent from the training set. These 150 compounds were arranged by predicted logP and vapour pressure values to provide a high-priority list of candidates for behavioral testing (Figure 3C).

### Predicted Natural Odors Strongly Repel *Drosophila* and mosquitoes

In order to test for repellency we first used *Drosophila melanogaster* in the 2-choice trap assay that previously showed DEET repellency(Reeder et al., 2001; Syed et al., 2011). We selected 4 affordable compounds from the list, (Methyl N,N-Dimethyl anthranilate, Ethyl anthranilate, Butyl anthranilate, 2,3-dimethyl-5-isobutyl pyrizine) the first 3 of which have been thoroughly tested for human use, have excellent safety profiles, a very mild and pleasant aroma like grapes, and have already been approved for human consumption/oral inhalation by the FDA, World Health Organization and European Food Safety Authority (EFSA, 2008; JECF, 2007) (Figure 3D). The pyrizine is a natural compound produced by ants as a trail pheromone (Tentschert et al., 2000). All 4 of the predicted compounds showed a very strong repellent effect on *Drosophila* across multiple doses tested using the trap-based 2-choice assay (Reeder et al., 2001; Syed et al., 2011) (Figure 3E).

One of the major disadvantages of DEET is its property of solubilizing plastics and synthetic materials (Krajick, 2006), which affects its usefulness by the military, and at home. We tested the ability of the 4 repellents to dissolve a 3 × 3mm square of vinyl. The plastic square disappeared completely in DEET within 6 hrs; however in the 4 predicted repellents there was no significant difference in the weight of the vinyl squares after 6 hrs, and even at the 30hr time point (Figure 4A).

**Figure 4.**
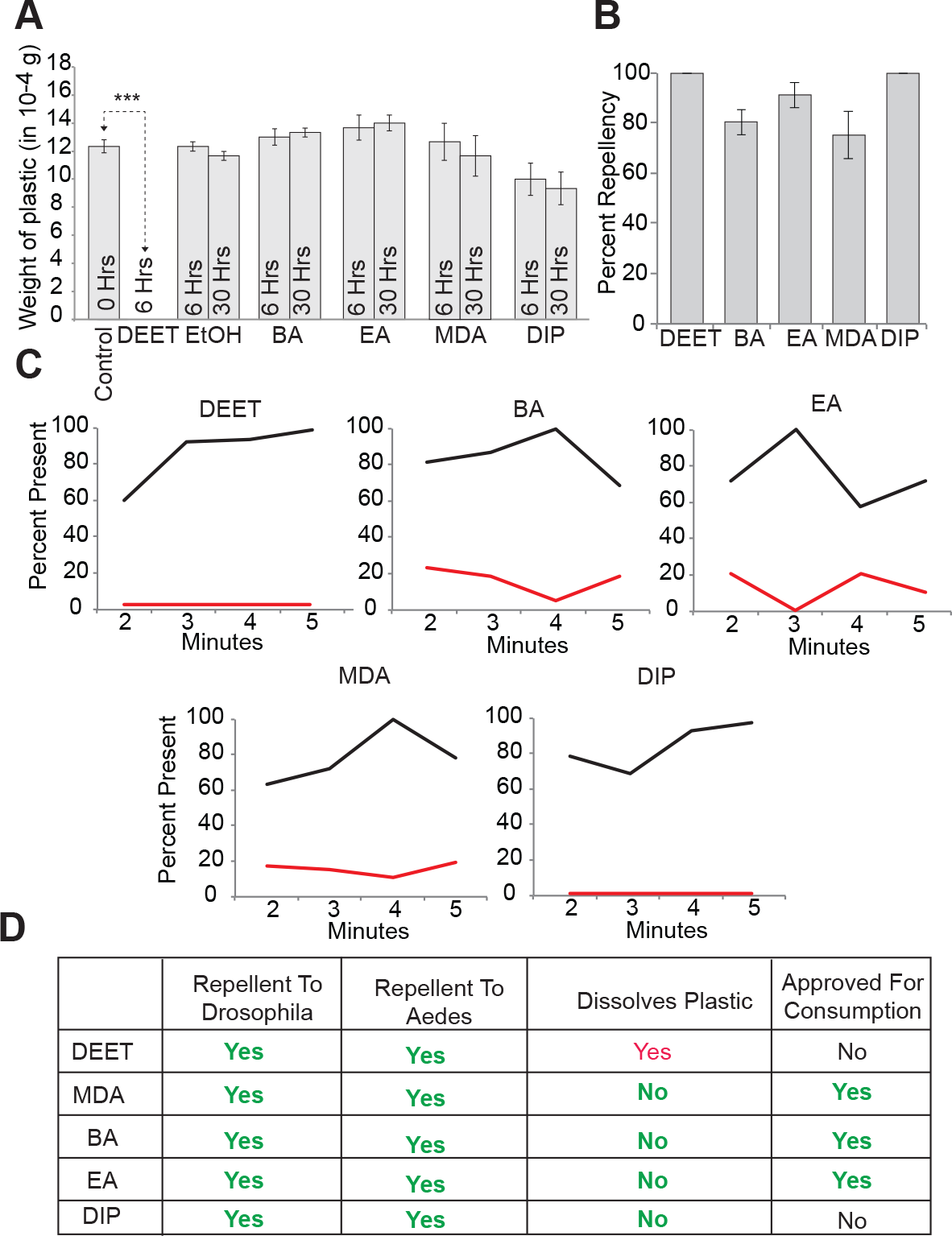
A new class of mosquito repellents with desirable safety profiles. **A,** Mean weight of vinyl pieces following submersion in indicated compounds for indicated amount of time. N=3, error bars=s.e.m., ***=p-value<10^−5^. **B,** Cumulative percentage of repellency across minutes 2,3,4 and 5 of indicated treatment (10%), in comparison to the appropriate solvent control. N=5 trials/treatment, 40 mosquitoes/trial. **C,** Mean percentage landing as measured by mosquitoes spending at least-5 secs on the protective window of glove, measured at different time-points in the 5-min assay. **D,** Summary of desirable properties of new insect repellents reported in this study.

In order to test whether these promising candidates were in fact repellent to mosquitoes, we performed behavior trials using the modified hand-in-a-cage assay. Notably, we find that all 4 compounds applied at 10% concentration demonstrate substantial repellency. Both the fraction of mosquitoes present on the net window over the arm throughout the duration of the assay, and the cumulative number of mosquitoes present on the net window were substantially lower in the test compounds as compared to solvent controls (Figures 4B and 4C). For the mosquitoes that did land on the repellent treatment, the escape index, as measured by the frequency of take off, was substantially higher as well (Figure S2).

### A New Generation of Safe and Effective Repellent Odors

Taken together we have identified a number of affordable and safe compounds with repellent properties, which establishes a significant advance toward the identification of DEET substitutes that are excellent candidates for regulatory approval for human and animal use (Figure 4D). This was made possible by a computational screening strategy that identified shared structural features of existing repellents to use for rapid identification of structurally related novel repellent compounds. Apart from the 4 compounds we have behaviorally identified as repellents, we have also used the screen to identify ~1000 novel compounds and more than a hundred additional natural compounds, many approved for use in human food and cosmetics, that may lead to several additional effective repellents for deadly insects. The repellency strategy can also have great potential if used in combination with other behavior control strategies such as masking of CO_2_-mediated attraction behavior or population control by trapping as a part of an integrated pull-mask-push approach(Turner et al., 2011; Turner and Ray, 2009). Since several of these compounds are affordable, repellent for fruit flies and approved for human consumption, they may have implications for control of agricultural pest insects as well that cause billions of dollars in crop loss as well as in protecting animals and pets that have tendencies to lick their skin. Moreover such substitutes may also have implications for control of DEET-resistant insects. We expect such repellents that are safe, affordable and do not solubilize plastics to pave the way for formulations that can be used in control of insect-human contact in disease endemic areas of the world and to provide an important line of defense against deadly diseases.

## EXPERIMENTAL PROCEDURES

### Behavior testing

Repellency was tested using *Drosophila melanogaster* 2-choice trap assay as described previously (Reeder et al., 2001; Syed et al., 2011).

*Preference Index*= (number of flies in treated trap − number of flies in control traps) /(number of flies in treated + control traps).

The *Drosophila* T-maze assay was conducted as described previously (Turner et al., 2011; Turner and Ray, 2009).

*Preference index* = (number of flies in test arm − number of flies in control arm)/(number of flies in test arm + number of flies in control arm).

Repellency was tested in mated and starved *Ae. aegypti* females using a hand-in-glove assay. Briefly, a gloved hand with an opening exposing skin odorants protected by 2 layers of netting was presented to mosquitoes for 5 min inside a cage and video taped for landing and avoidance responses. Mosquitoes were unable to bite due to the outer protective layer of netting and the inner layer of netting was treated with either test compound (10%) or solvent, such that mosquitoes were able to respond to volatiles but unable to make physical contact. The number of mosquitoes present for more than 5 seconds, and the numbers departing during the same period were counted from the videos at minutes 2,3,4, and 5 mins and repellency percentage and escape index calculated by comparing with similar numbers in solvent treated controls.

*Percentage repellency* = 100 × [1− (mean cumulative number of mosquitoes on the window of treatment for 5 seconds at time points 2,3,4,5 min/ mean cumulative number of mosquitoes that remained on window of solvent treatment for 5 seconds at time points 2,3,4,5 min)].

*Percentage present* = average number of mosquitoes on window for 5 seconds at a given time-point across trials. All values were normalized to percentage of the highest value for the comparison, which was assigned a 100 percent present.

*Mean Escape Index* = (Average Number of mosquitoes in treatment that landed yet left the mesh during a five second window over the following time points: 2 minutes, 3 minutes, 4 minutes, 5 minutes) / (Average Number of mosquitoes that landed yet left the mesh during a five second window over the same time points in (treatment + control)).

Each time point has N=5 trials, 40 mosquitoes per trial, Except for EA, where N=4.

### Chemical Informatics

Using a Sequential Forward Selection (SFS) approach 18 molecular descriptors from the 3,224 available from the Dragon package (Talete) were selected for their ability to increase the correlation between descriptor values and repellency. The Tune.SVM function from the R package was trained with the 18 descriptors for the training compound set using regression and a radial basis function kernel. We performed 20 independent 5-fold cross-validations followed by a receiver operating characteristics (ROC) analysis to analyze the performance of the computational repellency prediction method. The trained SVM then ranked both a library obtained from eMolecules (http://www.emolecules.com) (>400,000) and natural odor libraries (Supplementary Material) (>3,000)by repellency using the optimized descriptor set to identify the best candidate repellent compounds. The EPI Suite (http://www.epa.gov/oppt/exposure/pubs/episuite.htm) was used to calculate predicted LogP and Vapour Pressure values.

### Vinyl solubility test

One 3 × 3 mm square of 4-gauge vinyl was submerged in 1mL of each test compound in a glass container and stirred at a constant rate and weight measurements taken on a fine analytical balance at specified time-points.

### Supplemental Data

Supplemental Data include three figures and details of experimental procedures.

## ACKNOWLEDGMENTS

We thank I. Gomez, and Z. Wisotsky for comments on the manuscript.

